# Microsecond Interaural Time Difference Discrimination Restored by Cochlear Implants After Neonatal Deafness

**DOI:** 10.1101/498105

**Authors:** Nicole Rosskothen-Kuhl, Alexa N Buck, Kongyan Li, Jan W H Schnupp

**Affiliations:** Department of Biomedical Sciences, City University of Hong Kong, Hong Kong (SAR China); Neurobiological Research Laboratory, Section for Clinical and Experimental Otology, University Medical Center Freiburg, Freiburg, Germany

**Keywords:** Deafness, prosthetics, cochlear implant, binaural hearing, interaural time difference, psychoacoustics, hearing experience, inferior colliculus

## Abstract

Cochlear implants (CIs) can restore a high degree of functional hearing in deaf patients however spatial hearing remains poor, with many early deaf CI users reported to have no measurable sensitivity to interaural time differences (ITDs) at all. Deprivation of binaural experience during an early critical period is often blamed for this shortcoming. However, we show that neonatally deafened rats provided with precisely synchronized CI stimulation in adulthood can be trained to localize ITDs with essentially normal behavioral thresholds near 50 μs. Furthermore, neonatally deaf rats show high physiological sensitivity to ITDs immediately after binaural implantation in adulthood. The fact that our neonatally deaf CI rats achieved very good behavioral ITD thresholds while prelingually deaf human CI patients usually fail to develop a useful sensitivity to ITD raises urgent questions about whether shortcomings in technology or treatment may be behind the usually poor binaural outcomes for current binaural CI patients.

---

For people with severe to profound sensorineural hearing loss, cochlear implants (CIs) can be enormously beneficial routinely allowing near normal spoken language acquisition particularly when CI implantation takes place early in life [1]. Nevertheless the performance of CI users remains variable, and falls short of natural hearing.

Good speech understanding with competing sound sources requires the ability to separate speech from background. This is aided by “spatial release from masking”, which relies on the brain’s ability to process binaural spatial cues, including interaural level and time differences (ILDs & ITDs) [2]. Bilateral cochlear implantation is becoming commonplace to provide benefits of binaural hearing to the deaf [3-5], but binaural CI recipients perform poorly in sound localization and auditory scene analysis tasks, particularly when multiple sound sources are present [6,7]. The parameters that would allow CI patients to derive maximum benefits from binaural spatial cues are only partially understood. A number of technical problems (see [8], chapter 8) limit the fidelity with which CIs can encode ITDs in particular. Contemporary CI speech processors were originally designed for monaural, not binaural, hearing, and this likely contributes to the observed deficits in bilateral CI users [3]. Standard CI processors provide pulsatile stimulation which is not locked to the temporal fine structure of incoming sounds, and the timing of electrical pulses is not synchronized between both ears, making these devices incapable of encoding the temporal fine structure and ITDs with the required resolution of a few tens of μs. While normal hearing human listeners may be able to detect ITDs as small as 10 μs [9], ITD sensitivity of CI patients is poor or completely missing [3,5-7,10-12].

Whether CI patients can detect ITDs at all depends largely on their history: Prelingually deaf CI users invariably exhibit poor or no ITD sensitivity at all (rare star performers at best achieving thresholds of a few hundred µs), but many postlingually deaf CI users show at least some degree of ITD sensitivity [4,5,11-15]. This led to the suggestion that early auditory deprivation during a sensitive period may prevent the development of ITD sensitivity irrecoverably [1,15,16]. If this hypothesis is correct, then developing CI processors with better ITD coding might not benefit patients with hearing loss early in life because, by the time of implantation these patients would already have missed out on the formative sensory input needed to develop the brain circuitry required for binaural processing with microsecond precision. Various lines of evidence make this critical period hypothesis plausible, including immunohistochemical studies which have shown degraded tonotopic organization [17,18] and changes in stimulation-induced molecular, morphological, and electrophysiological properties of the auditory pathway of neonatally deafened (ND) CI-rats [17-21]. Additional studies demonstrate that abnormal sensory input during early development can alter ITD tuning curves in key brainstem nuclei of gerbils [22,23].

However, the “critical period hypothesis” of poor ITD sensitivity in CI patients has not been rigorously tested, and it is possible that the ND auditory pathway could retain an innate ability to encode ITD, which may then only be lost as a result of maladaptive plasticity after a period of binaural CI stimulation which conveys no useful ITD information. These possibilities cannot be distinguished based on clinical data, as there are no binaural CI processors in clinical use capable of encoding microsecond ITDs effectively. Thus, animal experimentation is needed, in a first instance, to examine how much functional ITD sensitivity can be achieved through bilaterally synchronized CI stimulation in a mature ND animal. Previous animal studies investigating binaural sensitivity after early deafness have been limited to acute electrophysiological experiments on cats and rabbits. These studies have reported reduced ITD sensitivity in the inferior colliculus (IC) [24-27] and auditory cortex (AC) [28,29] compared to hearing experienced animals. There have been no previous attempts to train ND animals to discriminate sub-millisecond ITDs across the physiological range delivered with CIs fitted in adulthood. Here, we show that cohorts of ND rats which received synchronized bilateral CI stimulation in young adulthood not only exhibit a great deal of physiological ITD sensitivity in their IC straight after implantation but they can also be trained easily and quickly to lateralize ITDs behaviorally, achieving thresholds as low as ∼50 μs, comparable to their normal hearing (NH) litter mates.

## Results

### Early deaf CI rats discriminate ITD as accurately as their normally hearing litter mates

To test whether ND rats can learn to discriminate ITDs of CI stimuli, we trained five ND rats who received chronic bilateral CIs in young adulthood (postnatal weeks 10-14) in a simple two-alternative forced choice (2AFC) ITD lateralization task, and we compared their performance against behavioral data from five age-matched NH rats trained to discriminate the ITDs of acoustic pulse trains [30]. Animals initiated trials by licking a center “start spout”, and responded to 200 ms long 50 Hz binaural pulse trains by licking either a left or a right “response spout” to receive drinking water as positive reinforcement (Figs. S2a, S3b). Which response spout would give water was indicated by the ITD of the stimulus. We used pulses of identical amplitude in each ear, so that systematic ITD differences were the only reliable cue available (Figs. S2c-d; S3f). ND rats were stimulated with biphasic electrical pulse trains delivered through chronic CIs, NH rats received acoustic pulse trains through near-field sound tubes positioned next to each ear when the animal was at the start spout (Fig. S3a). During testing, stimulus ITDs varied randomly. The behavioural data (Fig. 1) were collected over a testing period of around 14 days. For CI rats, the initial lateralization training started usually one day after CI implantation. On average, rats were trained for maximum eight days before we started to test them on ITD sensitivity. The behavioral performance of each rat is shown in Figure 1, using light blue for NH (a-e) and dark blue for ND (f-j) animals. Figure 1 clearly demonstrates that all rats, whether NH or ND with CIs, were capable of lateralizing ITDs. As might be expected, the behavioral sensitivity varied from animal to animal. To quantify the behavioral ITD sensitivity of each rat we fitted psychometric curves (see Methods, red lines in Fig. 1) to the raw data and calculated the slope of that curve at ITD=0. Figure 1k summarizes these slopes for NH (light blue) and ND CI (dark blue) animals. The slopes for both groups fell within the same range. Remarkably, the observed mean sensitivity for the ND animals (0.487 %/µs) is only about 20% worse than that of the NH (0.657 %/µs). Furthermore, the differences in means between experimental groups (0.17 %/µs) were so much smaller than the animal-to-animal variance (∼0.73 %^2^/µs^2^) that prohibitively large cohorts of animals would be required to have any reasonable prospect of finding a significant difference. Similarly, both cohorts showed similar 75% correct lateralization thresholds (median NH: 41.5 µs; ND: 54.8 µs; mean NH: 79.9 µs; ND: 63.5 µs). Remarkably, the ITD thresholds of our ND CI rats are thus orders of magnitude better than those reported for prelingually deaf human CI patients, who usually have ITD thresholds too large to measure, often in excess of 3000 µs [5,31]. Indeed, their thresholds are not dissimilar from the approx. 10-60 µs range of 75% correct ITD discrimination thresholds reported for normal human subjects tested with noise bursts [32], and pure tones [9], or the ≈ 40 µs thresholds reported for normally hearing ferrets tested with noise bursts [33].

**Figure 1:**
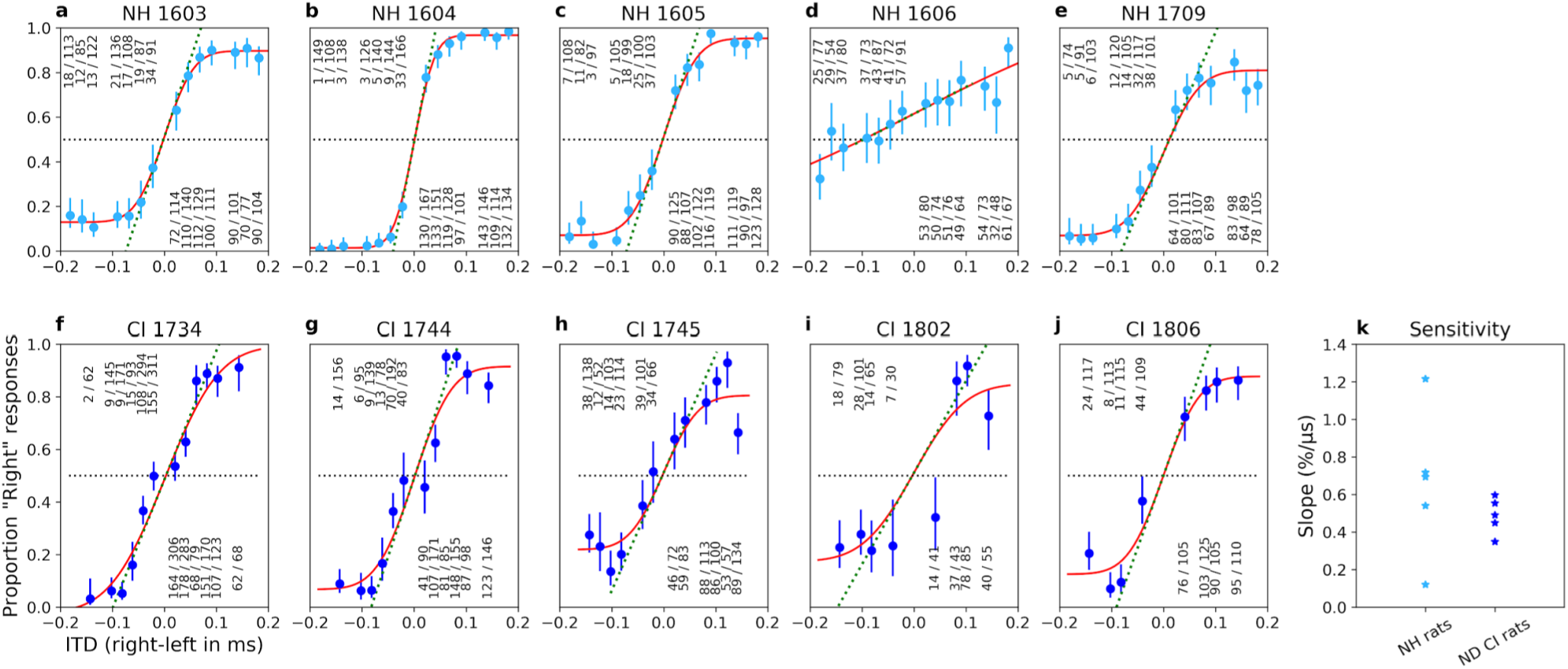
Early deaf CI rats discriminate ITD as accurately as their normally hearing litter mates. **a-j** ITD psychometric curves of normal hearing (**a-e**) and neonatally deafened CI rats (**f-j**). Panel titles show corresponding animal IDs. Y-axis: proportion of responses to the right-hand side. X-axis: Stimulus ITD in ms, with negative values indicating left ear leading. Blue dots: observed proportions of “right” responses for the stimulus ITD given by the x-coordinate. Number fractions shown above or below each dot indicate the absolute number of trials and “right” responses for corresponding ITDs. Blue error bars show Wilson score 95% confidence intervals for the underlying proportion “right” judgments. Red lines show sigmoid psychometric curves fitted to the blue data using maximum likelihood. Green dashed lines show slopes of psychometric curves at x=0. Slopes serve to quantify the behavioral sensitivity of the animal to ITD. Panel **k** summarizes the ITD sensitivities (psychometric slopes) across the individual animal data shown in **a-j** in units of % change in animal’s’ “right” judgments per μs change in ITD.

### Varying degrees and types of ITD tuning are pervasive in the neural responses in the IC of ND rats immediately after adult cochlear implantation

To investigate the amount of physiological ITD sensitivity present in the hearing inexperienced rat brain, we recorded responses of n=1140 multi-units in the IC of four young adult ND rats (Fig. 2, left) to isolated, bilateral CI pulse stimuli with ITDs varying randomly over a ±160 μs range (ca 123% of the rat’s physiological range [34]). For comparison, we also recorded responses of n=1312 IC multi-units in four age-matched NH rats (Fig. 2, right), using identical stimulation. For both cohorts, the stimuli were again biphasic current pulses of identical amplitude in each ear, so that systematic differences in responses can only be attributed to ITD sensitivity (see Fig. S2c-d). Responses of IC neurons were detected for currents as low as 100 μA. Figure 2 shows a selection of responses as raster plots and corresponding ITD tuning curves for both cohorts (Fig. 2, #1-8).

**Figure 2:**
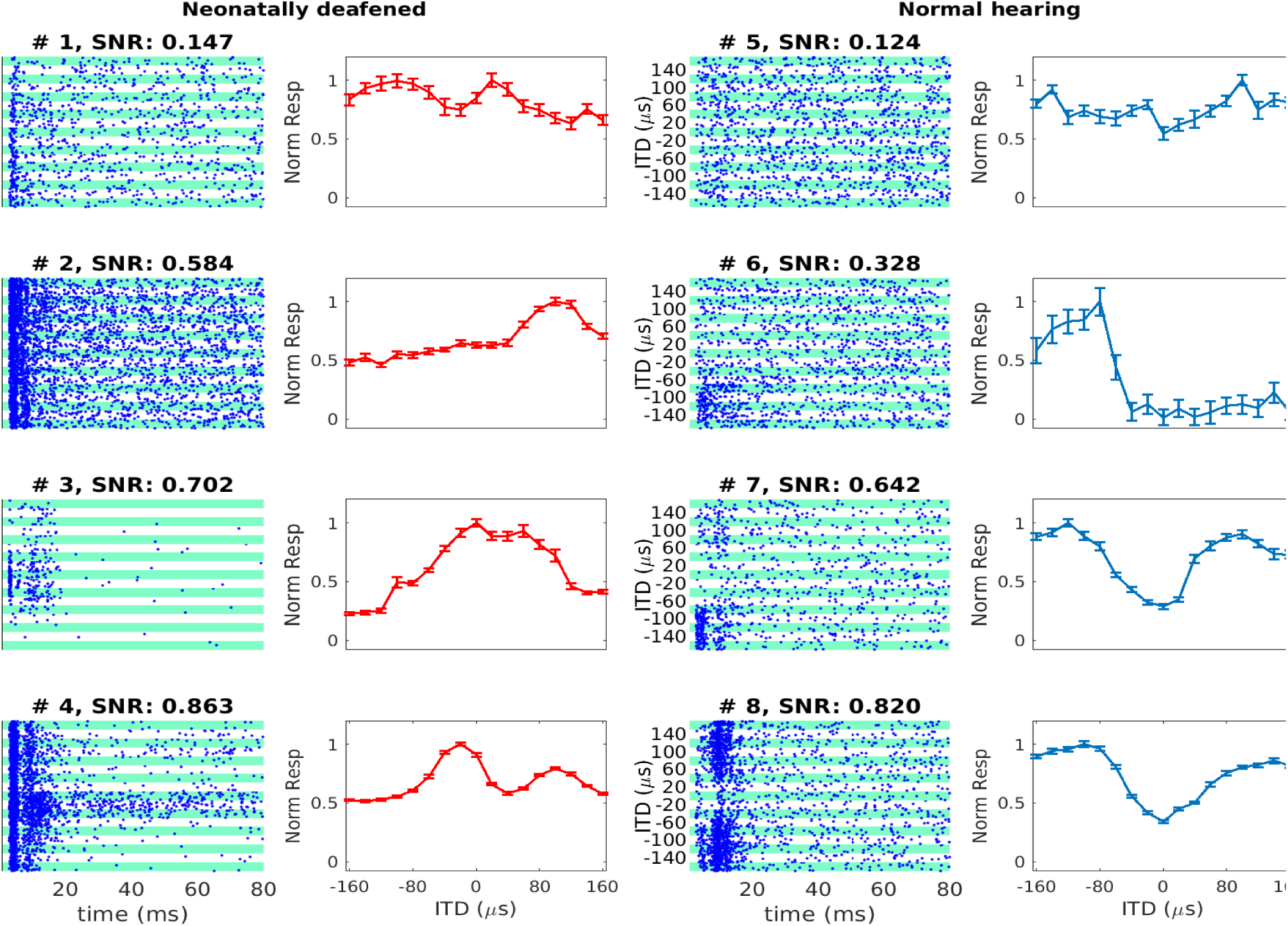
IC neurons of ND rats were ITD sensitive immediately following adult cochlear implantation. Examples of ITD tuning curves of neonatally deafened (ND) rats (red, left) and normal hearing (NH) rats (blue, right) with varying degrees and types of ITD sensitivity. Dot raster plots are shown next to corresponding ITD tuning curves. The multi-units shown were selected to illustrate some of the variety of ITD tuning curve depths and types observed. In the raster plots, each blue dot shows one spike. Alternating white and green bands show responses to n=30 repeats at each ITD shown. Tuning curve response amplitudes are baseline corrected and normalized relative to the maximum of the mean response across all trials, during a period of 3-80 ms post stimulus onset. Error bars show SEM. Above each sub-panel we show signal-to-noise (SNR) values to quantify ITD tuning. Panels are arranged top to bottom by increasing SNR. ITD>0: ipsilateral ear leading; ITD<0: contralateral ear leading.

Both for the ND and the NH animals, the manner in which varying ITD changed neural discharge patterns varied from one recording site to the next. But the extent to which discharge patterns depend on stimulus ITD is not obviously very different in the ND and the NH examples shown. While many multi-units showed typical short-latency onset responses to the stimulus with varying response amplitudes (Fig. 2, #1, #3, #6, #7), some showed sustained, but still clearly tuned, responses extending for up to 80 ms or longer post-stimulus (Fig. 2, #4). The shapes of ITD tuning curves we observed in rat IC (Fig. 2) resembled mostly the “peak”, “monotonic sigmoid”, “trough”, and “multi-peak” shapes previously described in the IC of cats [35].

To quantify how strongly the neural responses recorded at any one site depended on stimulus ITD, a “signal-to-noise ratio” (SNR) value was calculated as described in [24]. It quantifies the proportion of response variance that can be accounted for by stimulus ITD (see Methods). Each sub-panel of Figure 2 indicates SNR values obtained for the corresponding multi-unit, while Figure 3 shows the distributions of SNR values for both ND (dark red) and NH cohorts (light red). Figure 3 also shows the SNR values reported by Hancock et al. [24] for the IC of congenitally deaf (dark blue) and hearing experienced (light blue) cats. When comparing the distributions shown, it is important to be aware that there are significant methodological, as well as species, differences between our study and the study that produced the cat data shown in [24], so the cross-species comparison in particular must be done with care. Nevertheless, the distributions clearly show that ITD SNRs in our ND and NH rats are broadly similar, and also broadly in line with the values reported by others using similar methodologies. It is noticeable, and perhaps surprising, that the proportion of multi-units with relatively large SNRs values (substantial ITD tuning) is if anything slightly higher among the ND rats compared to the NH rats, and the median SNR value for ND rat IC multi-units (0.37) was in fact larger than that observed in NH rats (0.25). The proportion of rat multi-units which showed statistically significant ITD tuning (p≤0.01), as determined by ANOVA (see Methods), was also similarly very large in both ND (1041/1140 ≈ 91%) and NH CI-stimulated rats (1154/1312 ≈ 88%). Thus, for our rats which were kanamycin deafened before the onset of hearing, a lack of early auditory experience did not produce a measurable decline in overall sensitivity of IC neurons to the ITD of CI stimulus pulses. This is perhaps unexpected given that previous studies comparing ITD tuning in the IC of congenitally deaf white cats and with that in wild type cats did report noticeably worse ITD tuning in the congenitally deaf cats [26].

**Figure 3:**
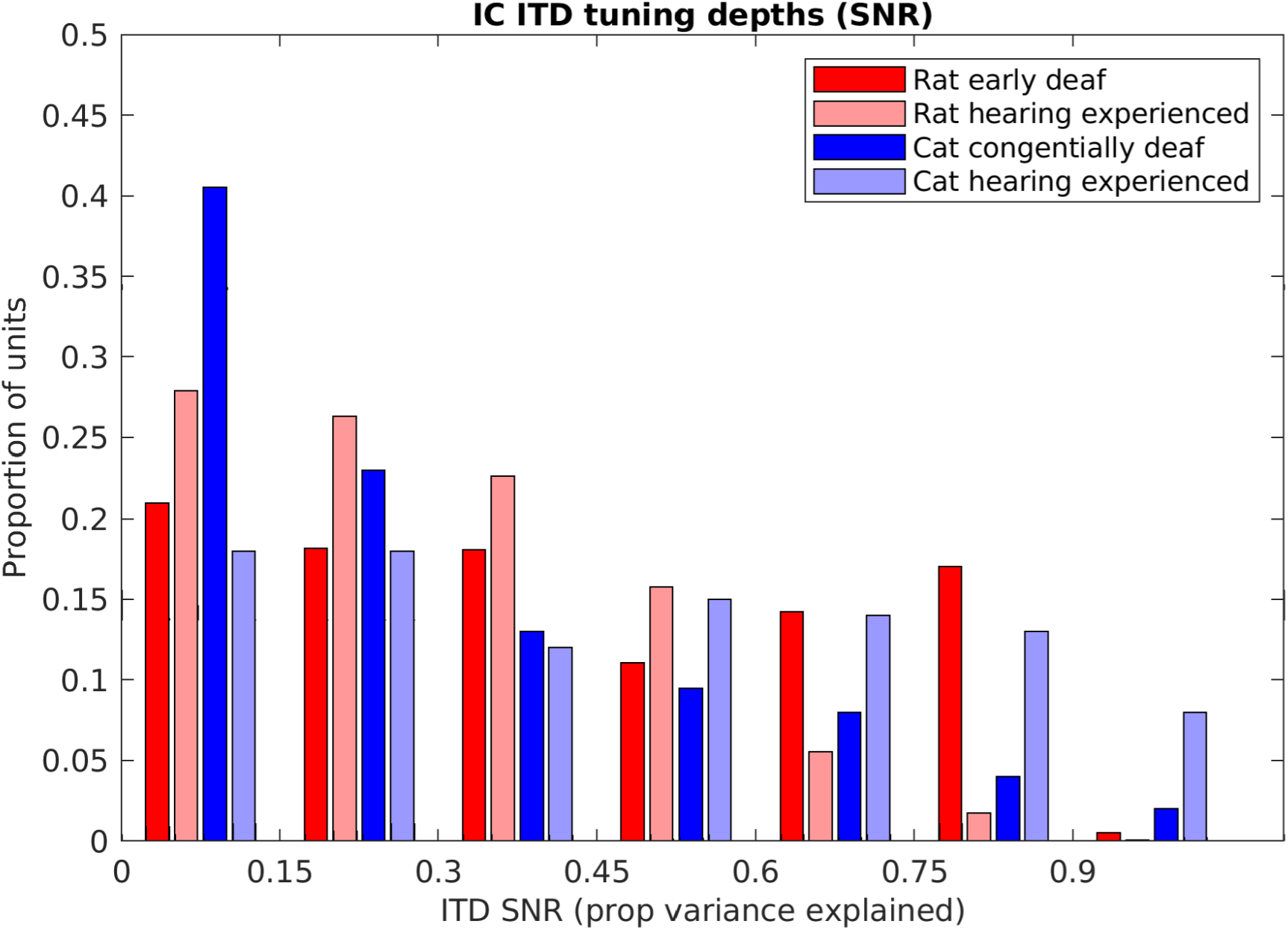
IC multi-units with good ITD tuning, as measured by high signal-to-noise ratio (SNR) values, are not less common in ND than in NH rats. Bar chart shows distributions of ITD SNR values for ICs multi-units of ND (red) and NH (light red) rats. SNR value distributions for IC single unit data recorded by Hancock et al. [24] for congenitally deaf (dark blue) and hearing experienced cats (light blue) are also shown for comparison.

Nevertheless, the results in Figures 2 and 3 clearly show that many IC neurons in the inexperienced, adult midbrain of ND rats are quite sensitive to changes in ITD of CI pulse stimuli by just a few tens of μs, and our behavioral experiments showed that ND rats (Fig. 1, f-j) can readily learn to use this neural sensitivity to perform behavioral ITD discrimination with an accuracy similar to that seen in their NH litter mates (Fig. 1, a-e).

## Discussion

This study is the first demonstration that, at least in rats, severely degraded auditory experience in early development does not inevitably lead to impaired binaural time processing in adulthood. In fact, the ITD thresholds of our ND rats (≈ 50 µs) were as good as the ITD thresholds of NH rats [30], and many times better than those typically reported for early deaf human CI patients with thresholds often too large to measure [5,14,30,31]. The surprisingly good performance exhibited by our ND animals raises the important question of whether early deaf human CI patients might perhaps also be able to achieve near normal ITD sensitivity if supplied with better binaural CI stimulation from the beginning of implantation. But before we consider translational questions that might be raised by our results, we should address two aspects of this study which colleagues may find surprising:

Firstly, some studies deemed rats to be generally poor at processing ITDs [36,37]. However, the only previous rat behavioral study outside of our lab tested interaural phase sensitivity only of relatively low frequency tones. We focused on broad-band acoustic or electrical pulse stimuli which provide plenty of “onset” and “envelope” ITDs, and which are processed well even at high carrier frequencies [38,39]. That may also explain why our CI rats showed good ITD sensitivity even though our CIs targeted the cochleas’ mid-frequency region, and not the apical region associated with low frequency hearing. Recent studies in CI patients with late deafness have shown that ITDs delivered to mid and high-frequency cochlear regions can be detected behaviorally [5,40].

Secondly, electrophysiological studies on congenitally deaf CI cats reported a substantially reduced ITD sensitivity relative to wild type, hearing experienced cats [24,28,29]. These studies recorded neural tuning high up in the auditory pathway (AC and IC), so one cannot be certain whether the reduced sensitivity reflects a fundamental degradation of ITD processing in the olivary nuclei, or merely poor maturation of connections to higher order areas, which may be reversed with experience and training. In the IC of our ND rats we found significant ITD sensitivity in 91% of recordings sites, compared to only 48% reported for congenitally deaf cats [24] or 62% for early deaf rabbits [27]. The proportion of ITD sensitive sites in ND rats is thus more similar to proportions in adult deafened CI-stimulated cats (84%-86%; [24,35], rabbits (∼75%; [27,41]) or gerbils (∼74%; [42]) and is comparable with our own NH rats (88%). Figure 3 suggests that the ITD SNR seen in our ND and NH CI-rats fall in a similar range of the ITD SNRs previously reported for congenitally deaf and hearing experienced CI-cats [24]. While for cats, the proportion of IC multi-units with large ITD SNR values appears to be reduced in animals lacking early auditory experience, the same does not appear to be the case in our rats. Whether these quantitative differences in physiological ITD sensitivity are due to methodological and/or species differences is not determinable, but we believe that these apparent differences are ultimately unlikely to be important, because even the congenitally deaf cats still have a decent number of IC units showing fairly large amounts of ITD sensitivity. In fact, more than 20% of the congenitally deaf cat IC units have SNR values of 0.5 or higher, which indicates really rather good ITD sensitivity. It is important to remember that it is essentially unknown how much ITD tuning in the IC or AC is necessary, or whether this is species specific, to make behavioral ITD discrimination thresholds of ≈ 50 µs possible, but multi-units such as #3 and #4 shown in Figure 2 change their firing rates substantially when ITDs change by only a few tens of microseconds, and these multi-units have SNRs that are not outside the range reported for congenital cats. This level of physiological sensitivity probably ought to be sufficient to enable quite good behavioral ITD discrimination performance if only these animals could be trained and tested on an appropriate task. Thus, in our opinion, any previously reported reductions in physiological ITD sensitivity seen in the IC [27,41] or AC [28,29] of early deaf animals simply does not seem nearly large enough to fully explain the extremely poor behavioral ITD thresholds seen in most early deaf humans and the observation that the physiological ITD sensitivity in our ND rats is comparable to that in NH animals [30] emphasizes the existence of other causes. Consequently, why in congenitally deaf cats there appears to be a relatively modest reduction in neural ITD sensitivity while in ND rats that appears not to be the case is not a pressing question, given that the available evidence suggests that both have quite a lot of innate or residual ITD sensitivity in their midbrains, and consequently ought to be able to learn to make use of that binaural cue. Thus, our finding of apparently normal behavioral ITD sensitivity in ND CI-rats may appear surprising, but it does not contradict previous studies [24,26-29].

The most striking difference between our results and other previously published work remains the vastly better behavioral ITD discrimination in ND CI rats compared to that of early deaf CI patients [5,14,15]. Previous authors have proposed that the very poor performance seen in these patients may be due to “factors such as auditory deprivation, in particular, lack of early exposure to consistent timing differences between the ears” [5]. However, a lack of early exposure did not prevent our ND rats from achieving very good ITD discrimination performance. Furthermore, our rats were neonatally deafened and implanted with CI in early adulthood thus not only lacking auditory experience during a so called ‘critical period’ but also having prolonged deprivation. This is comparable with prelingually deafened CI users who receive CIs as adults. However, Litovsky et al. [11] demonstrated that this cohort too had worse ITD performance as compared with adult-onset deafened CI users. Admittedly, there may be species differences at play here. Our ND rats were implanted as young adults, and were severely deprived of auditory input throughout their childhood, but humans mature much more slowly, so even patients implanted at a very young age will have suffered auditory deprivation for a substantially longer absolute time period than our rats. Nevertheless, our results raise the possibility that the poor ITD sensitivity of early deaf patients may perhaps not be caused by the “lack of exposure to consistent timing differences” and other factors might be to blame, such as the prolonged exposure to entirely inconsistent ITDs delivered by current clinical CI processors, which do not synchronize pulses between ears. This argument is supported by Zheng et al. [43] who found that CI implanted children with bilateral experience (>4 years) still show a higher percentage of errors than age-matched NH peers and Litovsky and Gordon [15] describe that “even the bilateral CI users who have had >6 years of experience listening with their CIs did not perform as well as the children with NH”. Additionally, most binaural patients receive their CIs sequentially, and their initial, potentially formative, auditory experience is monaural. In contrast, our rats received highly precise and informative ITDs right from the start.

Developmental studies in ferrets have shown that the formation of afferent synapses to medial superior olive, one of the main brainstem nuclei for ITD processing, is essentially complete before the onset of hearing [44]. In mice, the highly specialized calyx of held synapses which are thought to play key roles in relaying precisely timed information in the binaural circuitry have been shown to mature before the onset of hearing [45]. In gerbils, key parts of the binaural ITD processing circuitry in the auditory brainstem will fail to mature when driven with strong, uninformative omnidirectional white noise stimulation during a critical period [22,33,46-48], but no studies have demonstrated that critical periods in the ITD pathways will irrevocably close if sensory input is simply absent. In fact Tirko and Ryugo [49] show that inhibitory pathways in the medial superior olive, which are thought to be essential for ITD encoding [50], are significantly reduced in congenitally deaf cats at postnatal day 90, compared to normal hearing peers, and can be fully restored with the advent of CI stimulation after only 3 months. These data are therefore not incompatible with our hypothesis that inappropriate input, rather than its absence, may cause the loss of ITD sensitivity in early deaf CI users.

It is well known that the normal auditory system not only combines ITD information with ILD and monaural spectral cues to localize sounds in space, it also adapts strongly to changes in these cues, and can re-weight them depending on their reliability [29,51-53]. Current CI processors produce pulsatile stimulation based on fixed rate interleaved sampling, which is neither synced to stimulus fine structure nor synchronized between ears. Consequently, they only ever provide uninformative ITDs to the children using them. Perhaps brainstem circuits of children fitted with conventional binaural CIs simply “learn” to ignore unhelpful inputs which would be adaptive to them given that no useful information comes from the ITDs received. In contrast, precise ITD cues were essentially the only form of useful auditory input that our ND CI rats experienced, and they quickly learned to use them. Thus, our data raise the possibility that the mammalian auditory system may develop ITD sensitivity in the absence of early sensory input, and that this sensitivity is then either refined or lost depending on how informative binaural input turns out to be. The inability of early deaf CI patients to use ITDs may thus be somewhat similar to conditions such as amblyopia or failures of stereoscopic depth vision development, pathologies which are caused more by unbalanced or inappropriate inputs than by a lack of sensory experience [54]. For the visual system, it has been shown that orientation selective neuronal responses exist at eye-opening and thus are established without visual input [55]. If this hypothesis is correct, then it may be possible to “protect” ITD sensitivity in young binaural CI users by exposing them to regular periods of precise ITD information from the beginning of binaural stimulation. Whether CI patients are able to recover normal ITD sensitivity much later if rehabilitated with useful ITDs for prolonged periods, or whether their ability to process microsecond ITDs atrophies irreversibly, is unknown.

While the interpretations of our findings would lead us to argue that binaural CI processing strategies may need to change to make microsecond ITD information available to CI patients, one must nevertheless acknowledge the difficulty in changing established CI processing strategies. The continuous interleaved sampling (CIS) paradigm [56] from which most processor algorithms are derived, times the stimulus pulses so that only one electrode channel delivers a pulse at any one time. This is thought to minimize cross-channel interactions due to “current spread” which might compromise the already quite limited tonotopic place coding of CIs. Additionally, CI processors run at high pulse rates (≥900 Hz), which seems necessary to encode sufficient amplitude modulations (AM) for speech recognition [57]. However, ITD discrimination deteriorates when pulse rates exceeded a few hundred Hz in humans [58,59] and animals [41]. This is likely related to the observation that the ability of superior olivary neurons to encode envelope ITDs declines at envelope rates exceeding several hundred hertz [60]. Our own behavioral experiments described here were conducted with low pulse rates (50 Hz), and future work will need to determine whether ITD discrimination performance would be as good at pulse rates close to 1 kHz. Thus, designers of novel binaural CI speech processors may face conflicting demands: They must invent devices which fire each of 20 or more electrode channels in turn, at rates that are both fast, so as to encode speech AM in fine detail, but also slow, so as not to overtax the brainstem’s ITD extraction mechanisms, and they must make the timing of at least some of these pulses encode stimulus fine structure and ITDs. While difficult, this may not be impossible, and promising research is underway which either uses a mixture of different pulse rates for different electrode channels [61], or aims to “reset” the brain’s ITD extraction mechanisms by introducing occasional “double pulses” into the stimulus [62]. A detailed discussion is beyond the scope of this paper. Our results underscore the need to pursue this work with urgency, as it might just save many thousands of deaf patients from a loss of ITD sensitivity that could be caused by sub-optimal binaural CI-stimulation.

## Material and Methods

All procedures involving experimental animals reported here were approved by the Department of Health of Hong Kong (#16-52 DH/HA&P/8/2/5) or Regierungspräsidium Freiburg (#35-9185.81/G-17/124), as well as by the appropriate local ethical review committee.

### Deafening

Rats were neonatally deafened by daily intraperitoneal (i.p.) injections of 400 mg/kg kanamycin from postnatal day 9 to 20 inclusively [17,63]. This is known to cause widespread death of inner and outer hair cells [63-65] while keeping the number of spiral ganglion cells comparable to that in untreated control rats [63,65]. Osako et al. [63] have shown that rat pups treated with this method achieve hearing thresholds around 70 dB for only a short period (∼p14-16) and are severely to profoundly hearing impaired thereafter, resulting in widespread disturbances in the histological development of their central auditory pathways, including a nearly complete loss of tonotopic organisation [17,18,21]. We verified that this procedure provoked profound hearing loss (> 90 dB) by the loss of Preyer’s reflex [66], and the absence of auditory brainstem responses (ABRs) to broadband click stimuli (Fig. S1b). ABRs were measured as described in [21]: under ketamine (80mg/kg) and xylazine (12 mg/kg) anesthesia each ear was stimulated separately through hollow ear bars with 0.5 ms broadband clicks with peak amplitudes up to 130 dB SPL delivered at a rate of 43 Hz. ABRs were recorded by averaging scalp potentials measured with subcutaneous needle electrodes between mastoids and the vertex of the rat’s head over 400 click presentations. While normal rats typically exhibited click ABR thresholds near 30 dB SPL (Fig. S1a), deafened rats had very high click thresholds of ≥130 dB SPL; Fig. S1b).

### CI implantation, stimulation and testing

All surgical procedures, including CI implantation and craniotomy, were performed under anesthesia induced with i.p. injection of ketamine (80mg/kg) and xylazine (12 mg/kg). For maintenance of anesthesia during electrophysiological recordings, a pump delivered an i.p. infusion of 0.9% saline solution of ketamine (17.8 mg/kg/h) and xylazine (2.7 mg/kg/h) at a rate of 3.1 ml/h. During surgical and experimental procedures the body temperature was maintained at 38°C using a feedback-controlled heating pad (RWD Life Sciences, Shenzhen, China). Further detailed descriptions of our cochlear implantation methods can be found in previous studies [17,67-70]. In short, two to four rings of an eight channel electrode carrier (ST08.45, Peira, Beerse, Belgium) were fully inserted through a cochleostomy in medio-dorsal direction into the middle turn of both cochleae. Electrically evoked ABRs (EABRs) were measured for each ear individually to verify that both CIs were successfully implanted and operated at acceptably low electrical stimulation thresholds, usually around 100 μA (Fig. S1c). EABR recording used isolated biphasic pulses (see below) with a 23 ms inter-pulse interval. EABR mean amplitudes were determined by averaging scalp potentials over 400 pulses for each stimulus amplitude. For electrophysiology experiments, EABRs were also measured immediately before and after IC recordings, and for the chronically implanted rats, EABRs were measured once a week under anesthesia to verify that the CIs functioned properly and stimulation thresholds were stable.

### Electric and acoustic stimuli

The electrical stimuli used to examine the animals’ EABRs, the physiological, and the behavioral ITD sensitivity were generated using a Tucker Davis Technology (TDT, Alachua, Florida, US) IZ2MH programmable constant current stimulator at a sample rate of 48,828.125 Hz. The most apical ring of the CI electrode served as stimulating electrode, the next ring as ground electrode. All electrical intracochlear stimulation used biphasic current pulses similar to those used in clinical devices (duty cycle: 61.44 µs positive, 40.96 µs at zero, 61.44 µs negative), with peak amplitudes of up to 300 μA, depending on physiological thresholds or informally assessed behavioral comfort levels (rats will scratch their ears frequently, startle or show other signs of discomfort if stimuli are too intense). For behavioral training we stimulated all CI rats 6 dB above these thresholds. For details on the calibration which confirmed that our CI setup delivered the desired ITDs and no useable intensity cues, see supplementary Figure S2. Acoustic stimuli used to measure behavioral ITD sensitivity in NH rats were single sample pulse clicks generated at a sample rate of 48,000 Hz via a Raspberry Pi 3 computer connected to a USB sound card (StarTech.com, Ontario Canada, part # ICUSBAUDIOMH), amplifier (Adafruit stereo 3.7W class D audio amplifier, New York City, US, part # 987) and miniature high fidelity headphone drivers (GQ-30783-000, Knowles, Itasca, Illinois, US) which were mounted on hollow tubes. Stimuli were delivered at sound intensities of ≈ 80 dB SPL. For details on the calibration of the acoustic setup delivered the desired ITDs and no useable intensity cues, see supplementary Figure S3 and [30].

To produce electric or acoustic stimuli of varying ITDs spanning the rat’s physiological range of ± 130 µs [34], stimulus pulses of identical shape and amplitude were presented to each ear, with the pulses in one ear delayed by an integer number of samples. Given the sample rates of the devices used, ITDs could thus be varied in steps of 20.48 µs for the electrical, and 20.83 µs for the acoustic stimuli. The physiological experiments described here used single pulse stimuli presented in isolation, while the behavior experiments used 200 ms long 50 Hz pulse trains.

### Animal psychoacoustic testing

We trained our rats on 2AFC sound lateralization tasks using methods similar to those described in [30,33,47,48]. The behavioral animals were put on a schedule with six days of testing, during which the rats obtained their drinking water as a positive reinforcer, followed by one day off, with *ad-lib* water. The evening before the next behavioral testing period, drinking water bottles were removed. During testing periods, the rats were given two sessions per day. Each session lasted 25-30 min, which typically took 150-200 trials during which ≈ 10 ml of water were consumed.

One of the walls of each behavior cage was fitted with three brass water spouts, mounted ≈ 6-7 cm from the floor and separated by ≈ 7.5 cm (Figs. S2a-b; S3a-c). We used one center “start spout” for initiating trials and one left and one right “response spout” for indicating whether the stimulus presented during the trial was perceived as lateralized to that side. Contact with the spouts was detected by capacitive touch detectors (Adafruit industries, New York City, US, part # 1362). Initiating a trial at the center spout triggered the release of a single drop of water through a solenoid valve. Correct lateralization triggered three drops of water as positive reinforcement. Incorrect responses triggered no water delivery but caused a 5-15 s timeout during which no new trial could be initiated.

Timeouts were also marked by a negative feedback sound for the NH rats. Given that CI stimulation can be experienced as strenuous and tiring by human patients [71], the ND CI rats received a flashing LED as an alternative negative feedback stimulus. After each correct trial a new ITD was chosen randomly from a set spanning ±160 μs in 25 µs steps, but after each incorrect trial the last stimulus was repeated in a “correction trial”. Correction trials prevent animals from developing idiosyncratic biases favoring one side [47,72], but since they could be answered correctly without attention to the stimuli by a simple “if you just made a mistake, change side” strategy, they are excluded from the final psychometric performance analysis.

The NH rats received their acoustic stimuli through stainless steel hollow ear tubes placed such that, when the animal was engaging the start spout, the tips of the tubes were located right next to each ear of the animal to allow near-field stimulation (Fig. S3a). The pulses resonated in the tubes, producing pulse-resonant sounds, resembling single-formant artificial vowels with a fundamental frequency corresponding to the click rate. Note that this mode of sound delivery is thus very much like that produced by “open” headphones, such as those commonly used in previous studies on binaural hearing in humans and animals, e.g. [33,73]. We used a 3D printed “rat acoustical manikin” with miniature microphones in the ear canals (Fig. S3c). It produced a channel separation between ears of ≥ 20dB at the lowest, fundamental frequency and around 40 dB overall. Further details on the acoustic setup and procedure are described in [30]. The ND CI rats received their auditory stimulation via bilateral CIs described above, connected to the TDT IZ2MH stimulator via a custom-made, head mounted connector and commutator, as described in [74].

### Multi-unit recording from IC

Anesthetized rats were head fixed in a stereotactic frame (RWD Life Sciences), craniotomies were performed bilaterally just anterior to lambda. The animal and stereotactic frame were positioned in a sound attenuating chamber, and a single-shaft, 32-channel silicon electrode array (ATLAS Neuroengineering, E32-50-S1-L6) was inserted stereotactically into the left or right IC through the overlying occipital cortex using a micromanipulator (RWD Life Sciences). Extracellular signals were sampled at a rate of 24.414 kHz with a TDT RZ2 with a NeuroDigitizer headstage and BrainWare software. Our recordings typically exhibited short response latencies (≈ 3-5 ms), which suggests that they may come predominantly from the central region of IC. Responses from non-lemniscal sub-nuclei of IC have been reported to have longer response latencies (≈ 20ms; [75]).

At each electrode site, we first measured neural rate/level functions, varying stimulation currents in each ear to verify that the recording sites contained neurons responsive to cochlear stimulation, and to estimate threshold stimulus amplitudes. Thresholds rarely varied substantially from one recording site to another in any one animal. We then measured ITD tuning curves by presenting single pulse binaural stimuli with equal amplitude in each ear, ≈ 10 dB above the contralateral ear threshold, in pseudo-random order. ITDs varied from 163.84 μs (8 samples) contralateral ear leading to 163.84 μs ipsilateral ear leading in 20.48 μs (one sample) steps. Each ITD value was presented 30 times at each recording site. The inter-stimulus interval was 500 ms. At the end of the recording session the animals were overdosed with pentobarbitone.

### Data analysis

To quantify the extracellular multi-unit responses we calculated the average activity for each stimulus over a response period (3-80 ms post stimulus onset) as well as baseline activity (300-500 ms after stimulus onset) at each electrode position. The first 2.5 ms post stimulus onset were dominated by electrical stimulus artifacts and were discarded. For display purposes of the raster plots in Figure 2 we extracted multi-unit spikes by simple threshold crossings of the bandpassed (300 Hz - 6 kHz) electrode signal with a threshold set at four standard deviation of the signal amplitude. To quantify responses for tuning curves, instead of counting spikes by threshold crossings we instead computed an analog measure of multi-unit activity (AMUA) amplitudes as described in [76]. The mean AMUA amplitude during the response and baseline periods was computed by bandpassing (300 Hz - 6 kHz), rectifying (taking the absolute value) and lowpassing (6 kHz) the electrode signal. This AMUA value thus measures the mean signal amplitude in the frequency range in which spikes have energy. As illustrated in Figure 1 of [76], this gives a less noisy measure of multi-unit neural activity than counting spikes by conventional threshold crossing measures because the later are subject to errors due to spike collisions, noise events, or small spikes sometimes reach threshold and sometimes not. The tuning curves shown in the panels of Figure 2 were measured using this AMUA measure. It is readily apparent that changes in the AMUA amplitudes track changes in spike density.

### Signal-to-noise ratio (SNR) calculation

SNR values are a measure of the strength of tuning of neural responses to ITD which we adopted from [24] to facilitate quantitative comparisons. The SNR is defined in [24] as the proportion of trial-to-trial variance in response amplitude explained by changes in ITD. The SNR is calculated by computing a one-way ANOVA of responses grouped by ITD value and dividing the total sum of squares by the group sum of squares. This yields values between 0 (no effect of ITD) and 1 (response amplitudes completely determined by ITD). P-values were also computed from the one-way ANOVA and p < 0.01 served as a criterion to determine whether the ITD tuning of a given multi-unit was deemed statistically significant. The number of responses for each ITD value was 30, yielding with a degree of freedom (df) for the ANOVA of 29.

### Psychometric curve fitting

In order to derive summary statistics that could serve as measures of ITD sensitivity from the thousands of trials performed by each animal we fitted psychometric models to the observed data. It is common practice in human psychophysics to fit performance data with cumulative Gaussian functions [77,78]. This practice is well motivated in signal detection theory, which assumes that the perceptual decisions made by the experimental subject are informed by sensory signals which are subject to multiple, additive, and hence approximately normally distributed sources of noise. When the sensory signals are very large relative to the inherent noise then the task is easy and the subject will make the appropriate choice with near certainty. For binaural cues closer to threshold, the probability of choosing “right” (*p*_*R*_) can be modeled by the function

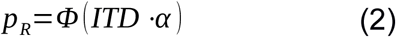

where, Φ is the cumulative normal distribution, *ITD* denotes the interaural time difference (arrival time at left ear minus arrival time at right ear, in ms), and *α* is a sensitivity scale parameter which captures how big a change in the proportion of “right” choices a given change in ITD can provoke, with units of 1/ms.

Functions of the type in equation (2) tend to fit psychometric data for 2AFC tests with human participants well, where subjects can be easily briefed and lack of clarity about the task, lapses of attention or strong biases in the perceptual choices are small enough to be explored. However, animals have to work out the task for themselves through trial and error, and may spend some proportion of trials on “exploratory guesses” rather than direct perceptual decisions. If we denote the proportion of trials during which the animal makes such guesses (the “lapse rate”) by *γ*, then the proportion of trials during which the animal’s responses are governed by processes which are well modeled by equation (2) is reduced to (1*-γ*). Furthermore, animals may exhibit two types of bias: an “ear bias” and a “spout bias”. An “ear-bias” exists if the animal hears the midline (50% right point) at ITD values which differ from zero by some small value *β*. A “spout bias” exists if the animal has an idiosyncratic preference for one of the two response spouts or the other, which may increase its probability of choosing the right spout by *δ* (where *δ* can be negative if the animal prefers the left spout). Assuming the effect of lapses, spout and ear bias to be additive, we therefore extended eqn (2) to the following psychometric model:

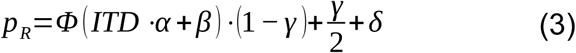

We fitted the model in equation (3) to the observed proportions of “right” responses as a function of stimulus ITD using the scipy.optimize.minimize() function of Python 3.4, using gradient descent methods to find maximum likelihood estimates for the parameters *α, β, γ* and *δ* given the data. This cumulative Gaussian model fitted the data very well, as is readily apparent in Figure 1a-j. We then used the slope of the psychometric function around zero ITD as our maximum likelihood estimate of the animal’s ITD sensitivity, as plotted in Figure 1k. That slope is easily calculated using the equation (4)

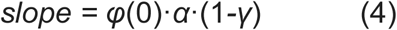

which is obtained by differentiating equation (3) and setting ITD=0. *φ(*0) is the Gaussian normal probability density at zero (≈0.3989).

Seventy-five % correct thresholds were computed as the mean absolute ITD at which the psychometric dips below 25% or rises above 75% “right” responses respectively.

## Acknowledgements

We thank A. Hyun Jung for assisting behavioral training of CI rats, P. Ruther and the *Cluster of Excellence BrainLinks-BrainTools* (German Research Foundation, grant number EXC1086) for the support by recording electrodes. Work leading to this publication was supported by a grant from the Vice President of Research and Technology (VPRT) of the City University of Hong Kong, the German Academic Exchange Service (DAAD) with funds from the German Federal Ministry of Education and Research (BMBF) and the People Programme (Marie Curie Actions) of the European Union’s Seventh Framework Programme (FP7/2007-2013) under REA grant agreement n° 605728 (P.R.I.M.E. – Postdoctoral Researchers International Mobility Experience), and friends’ association “Taube Kinder lernen hören e. V.”.

## Author Contributions

N.R.K. and J.W.H.S. designed the study and wrote the article. N.R.K., A.N.B., K.L., and J.W.H.S. performed the experiments. J.W.H.S., N.R.K., and A.N.B. evaluated the data. All authors approved the final manuscript.

## Declaration of Interests

The authors declare no competing interests.

## Supplementary Figures

**Figure S1:**
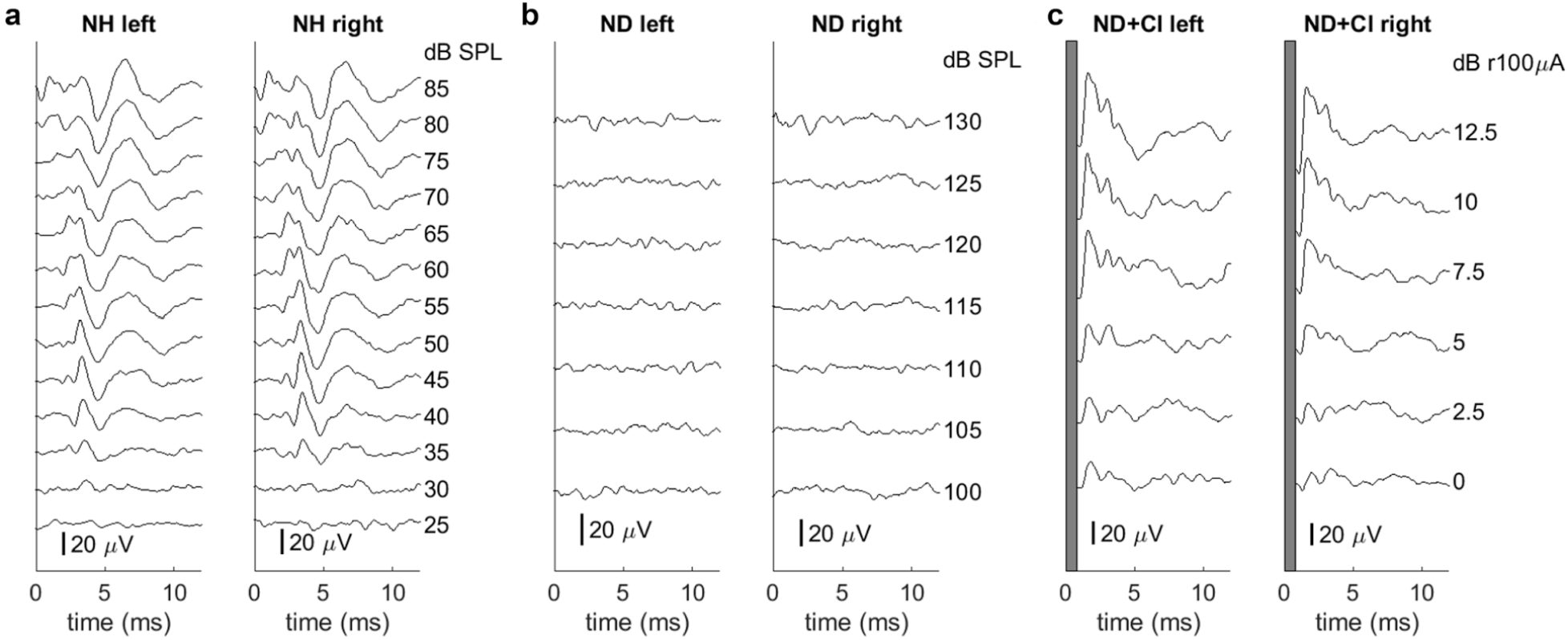
Brainstem recordings to verify normal hearing or loss of hearing function as well as the symmetrical placement of CIs. **a** Auditory brainstem responses of an acoustically stimulated normal hearing (NH) rat. ABRs are symmetrical for both ears and show clear differentiation. **b** ABRs of a neonatally deafened (ND) rat. No hearing thresholds were detectable up to 130 dB SPL. **c** Electrically evoked ABRs under CI stimulation of a deafened rat. Each sub-panel includes measurements for the left and the right ear, respectively, under acoustic (**a-b**) or electric stimulation (**c**). In (c) the first millisecond (electrical stimulus artifact) is blanked out.

**Figure S2:**
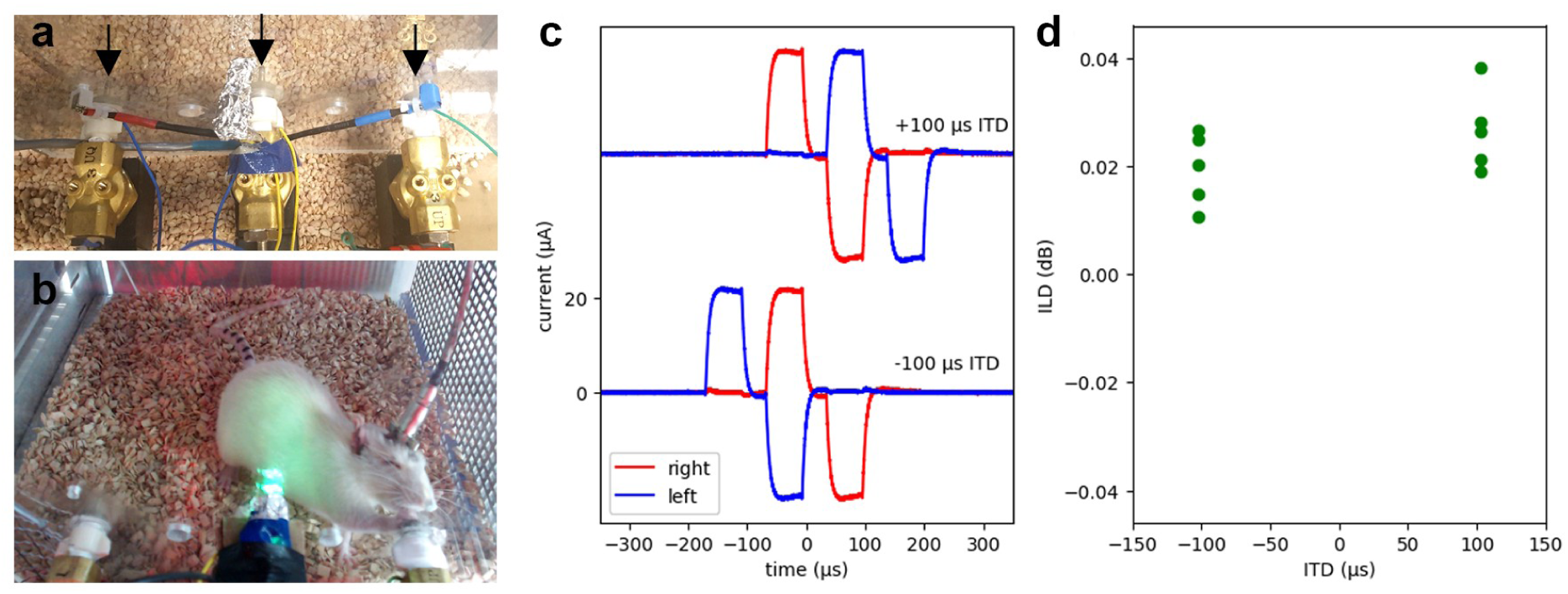
Binaural electrical intracochlear stimulation of CI rats. **a** Close-up of the training setup for CI rats. The central “start” and lateral “response” spouts deliver the water reward and are indicated by arrows. **b** CI rat during a testing session, making a response to the left by making contact with the left reward spout. **c** Calibration measurements were performed by connecting the stimulator cable to 10 kOhm resistors instead of the in-vivo electrodes and recording voltages using a Tektronix MSO 4034B oscilloscope with 350 MHz and 2.5 GS/s. Recordings of stimulus pulses are shown with 100 µs ITD leading in the right ear (top) or the left ear (bottom) respectively. Pulses delivered to the right ear are shown in red, those delivered in the left ear in blue. The stimulator was programmed to produce biphasic rectangular stimulus pulses with a 20 µA amplitude (y-axis) and a 20.5 µs interval between the positive and the negative phase. **d** Measured calibration pulses such as those shown in (**c**) were used to verify that electric ILDs were negligible and did not vary systematically with ITD. ILDs were computed as the difference in root mean square (RMS) power of the signals in (**d**). Data from five presentations of ITDs of + or – 100 µs are shown by the green dots. These residual ILDs produced by device tolerances in our system are not only an order of magnitude smaller than the ILD thresholds for human CI subjects reported in the literature (∼0.1 dB [79]), they also do not covary with ITD. We can therefore be certain that sensitivity to ILDs can not account for our behavior data.

**Figure S3:**
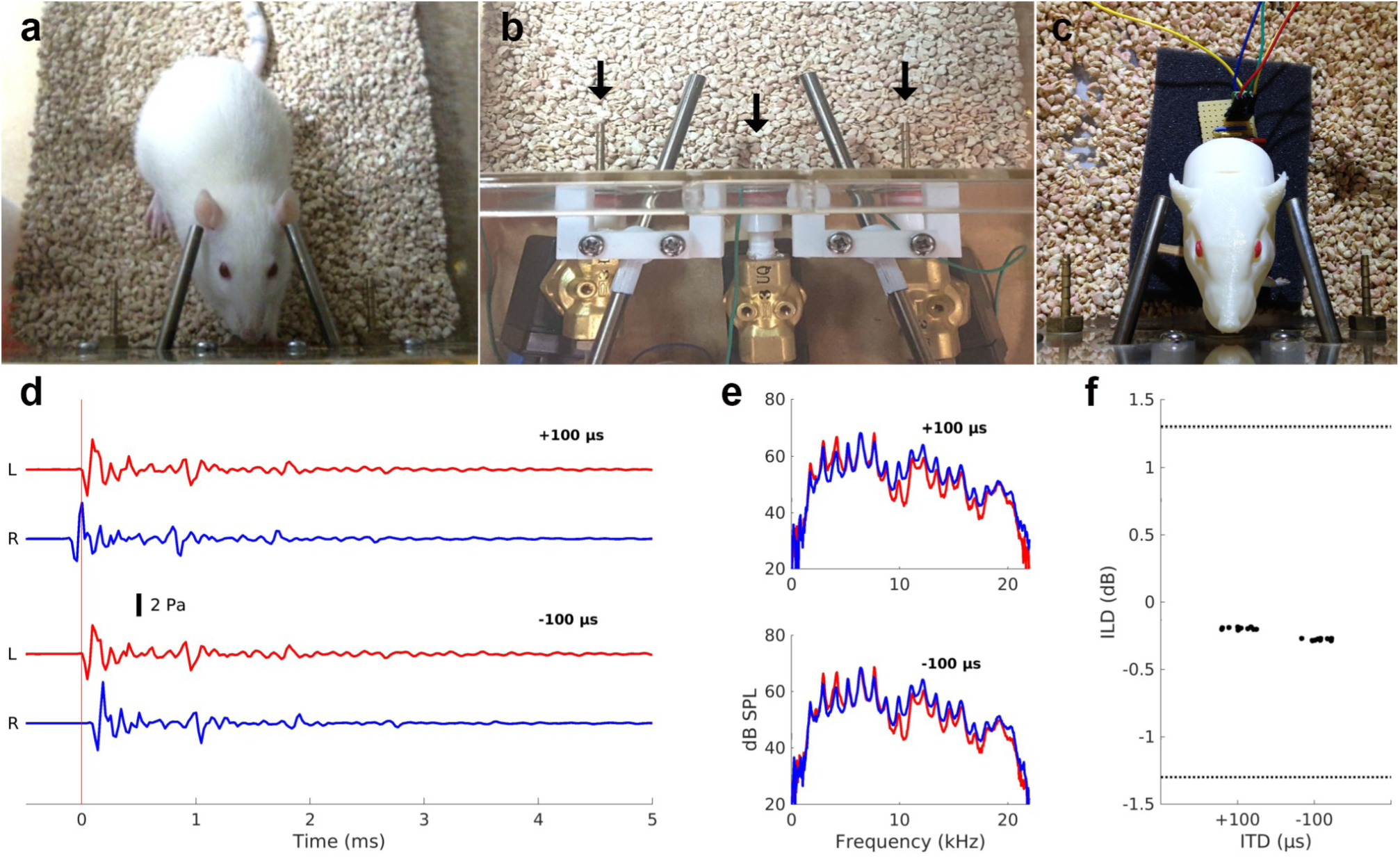
Binaural psychoacoustics near-field setup for NH rats. Note that the setup is identical to that described in Li et al. [30] and this supplementary figure is similar to Figure 1 of that paper. **a** NH rat during a testing session, initiating a trial by making contact with the central “start” spout. Steel tube phones are positioned close to each ear. **b** Close-up of the assembly. The central “start” and lateral “response” spouts deliver the water reward and are indicated by arrows. Also visible are the custom ball joints for adjusting the tube phone positions. **c** 3D printed “rat acoustical manikin” with miniature microphones in each ear canal, used for validating the setup. **d** Validation data for acoustic click stimuli as recorded from the microphones inside each ear canal of the 3D printed “rat acoustical manikin” (L: left ear, R: right ear) in response to the +/- 100 µs ITD conditions (top and bottom pair of traces, respectively). **e** Frequency spectra of the sound waveforms recorded by the microphones in each ear for the +100 µs (top) and -100 µs (bottom) conditions. **f** Acoustic ILDs (y-axes) measured through the “rat acoustical manikin” microphones for the +/- 100 µs ITD conditions. ILDs were computed as the difference in root mean square (RMS) power of the signals in panel (**d**). Data were recorded from 10 presentations of each ITD stimulus, and each dot represents one trial (a random amount of scatter along the x-axis was added for ease of visualization). Note that the residual ILDs are much smaller than the reported behavioral thresholds for ferrets (∼ 1.3 dB, dotted line, [33]) or rats (∼3 dB [37]). We can therefore be certain that sensitivity to ILDs cannot account for our behavior data.

## Notes

### Competing Interest Statement

The authors have declared no competing interest.

### Summary of Updates

text body revised

